# Enhanced iPSC Cardiomyocyte Maturation via Combined 3D-Culture and Metabolic Cues

**DOI:** 10.1101/2025.10.29.684362

**Authors:** Sabine Rebs, Hanna Eberl, Nicole Wagensohner, Nataliya Dybkova, Julia K. Unsöld, Jan Dudek, Patricia M. Costa, Marbely C. Fernandez, Eva A. Rog-Zielinska, Franziska Schneider-Warme, Christoph Maack, Samuel Sossalla, Katrin Streckfuss-Bömeke

**Affiliations:** Institute of Pharmacology and Toxicology, University of Würzburg, Germany; Clinic for Cardiology and Pneumology, University Medical Centre, Göttingen, Germany; Department of Translational Research, Comprehensive Heart Failure Centre (CHFC), University Clinic Würzburg, Germany; Institute for Experimental Cardiovascular Medicine, University Heart Center, Medical Center - University of Freiburg, and Faculty of Medicine, University of Freiburg, Germany; Medical Clinic I, Cardiology and Angiology, Justus-Liebig-University, Giessen, Germany; Department of Cardiology, Campus Kerckhoff, Bad Nauheim, Germany; Cardio-Pulmonary Institute (CPI), Giessen, Germany

## Abstract

Human induced pluripotent stem cell-derived cardiomyocytes (iPSC-CM) have become an invaluable tool for disease modelling and drug testing. However, while many etiologies of heart failure involve defects in excitation-contraction coupling, mitochondrial energetics or both, iPSC-CM are limited by the developmental immaturity of these processes. Here, we report a combinatorial strategy to enhance the maturation of human iPSC-CM by integrating three-dimensional (3D) spheroid culture conditions with a defined hormone- and fatty acid-enriched maturation medium (MM). A comprehensive analysis of structural, electrophysiological and Ca^2+^ handling parameters was performed to evaluate cellular and functional maturation. The iPSC-CM generated under these conditions (3D_MM) exhibit many phenotypic characteristics that resemble those of isolated adult human CM, including (i) a rod-shaped morphology, (ii) cardiac ultrastructural features such as aligned myofilaments, unidirectional organized sarcomeres, and the presence of transverse (t)-tubules, (iii) refined action potential (AP) parameters and Ca^2+^ handling, and (iv) β-adrenergic responsiveness and a positive force-frequency relationship. Compared with long-term (LT) monolayer cultures of 90 days or the individual cues (3D or MM alone), the 3D_MM protocol achieves mostly superior or at the least non-inferior maturation effects. This systematic investigation further demonstrates that while 3D culturing or MM alone improved specific aspects of maturation, only their synergistic combination produced a comprehensive enhancement of key CM processes, such as excitation-contraction coupling and mitochondrial energetics.

## Introduction

The heart is an efficient, continuously working engine that consumes vast amounts of energy, which must be replenished by oxidative phosphorylation of adenosine triphosphate (ATP) in mitochondria [1]. During exercise, cardiac output increases severalfold, and to match energy supply to demand, a tight coupling between excitation-contraction (EC) coupling and mitochondrial energetics is required, a process termed mechano-energetic coupling [2]. In various acquired, but also hereditary etiologies of heart failure, defects in EC coupling, mitochondrial energetics and/or mechano-energetic coupling can be the cause of disease or aggravate cardiac remodeling and dysfunction [2]. To model human physiology and pathology closely, human induced pluripotent stem cell-derived cardiomyocytes (iPSC-CM) have become an established and invaluable experimental platform in cardiovascular research and regenerative medicine. Unlike primary CM isolated from human myocardial tissue, which are rare to obtain and maintain in culture, iPSC-CM can be produced in large quantities from any donor, tailored to specific genetic backgrounds to generate isogenic, mutation-inserted, or reporter lines, and/or used as a source for cell-replacement therapy [3, 4]. However, a relevant limitation of iPSC-CM is that its immature state, as reflected in ultrastructural features, EC coupling, and mitochondrial metabolism, remains a significant constraint.

Postnatally, the heart undergoes further significant functional maturation. CM postnatal maturation is multifaceted and includes a metabolic shift from glycolysis to fatty acid oxidation (FOA), exit from the cell proliferation cycle and subsequent cell growth, and a transition towards binucleation. Furthermore, the processes of EC coupling become more efficient by the development of t-tubules that form dyads with the sarcoplasmic reticulum (SR) [5]. Freshly differentiated iPSC-CM cultured for 25-30 days without any further maturation cues resemble a more fetal developmental state of a CM, characterized by round morphology with a regular but shorter sarcomere length, lacking ultrastructural features of t-tubules or mitochondrial cristae, and exhibiting a lower oxidative capacity and longer action potentials (AP) with a more positive resting membrane potential (RMP), to name a few characteristic properties reflecting their immaturity [6-8].

iPSC-CM can faithfully recapitulate hereditary cardiomyopathies [9], while modeling of other, also acquired cardiac disease traits is more challenging or even obscured by the immaturity of iPSC-CM, which could potentially be unveiled by enhancing iPSC-CM maturation [10-12]. Also, the modelling of drug responsiveness depends on the maturation status. For example, immature iPSC-CM show lower sensitivity and efficacy to β-adrenergic stimulation than mature CM [13, 14].

Based on the growing demand to improve CM maturation protocols, significant progress has been made with the application of distinct maturation cues, such as optimized media composition, exposure to electrical or mechanical stimulation, cell seeding on nanopatterns, or introducing 3D culture formats [7]. A fully adult-like iPSC-CM phenotype, however, has not yet been achieved. Longterm (LT) culture is often utilized to foster maturation and represents an easy-to-implement, but time-consuming protocol applicable in most laboratories. Extended culture periods of at least 60 days promote structural and functional maturation. LT culture results in improved structural and sarcomeric organization, evident by formation of caveolae and increased sarcomeric regularity. Metabolically, an extended culture time enhances metabolic capacity with a shift towards FAO [15-17]. Moreover, LT culture positively affects electrophysiological function, resulting in higher L-type calcium (*I*_Ca,L_) and sodium (*I*_Na_) currents, shortened AP duration (APD) and an AP shape that more closely resembles adult cardiac AP [18]. Different hormone- and/or lipid-enriched maturation media (MM) also promote mitochondrial and metabolic maturation by enhancing oxidative capacity and shifting from glycolysis to FAO already at earlier timepoints than LT culture [19]. MM can also benefit cell shape and the expression of calcium (Ca^2+^) handling proteins, especially Ca^2+^ channels, and result in higher contractile force, but to a lesser extent than achieved by electrical or mechanical maturation protocols [19-21]. On one hand, electrical stimulation substantially matures iPSC-CM with respect to electrophysiological properties, promoting an adult-like AP shape, APD, and RMP. On the other hand, however, little is reported on metabolic effects of electrical stimulation [22, 23]. More technically sophisticated approaches include nanopatterning of iPSC-CM, mechanical stimulation (defined stretch-and-release), or casting iPSC-CM into engineered heart tissues. Although these approaches greatly boost maturity, especially on the structural and contractile level, they yet require specialized devices and technical skills for experimental implementation and analysis [13, 24-26].

Here, we comprehensively explore the synergetic potential of a novel-composite MM combined with a 3D-spheroid model to enhance iPSC-CM maturation that does not require any specialized technical equipment beyond standard cell culture materials. These insights advance our understanding of the maturation of ultrastructural features, EC coupling, mitochondrial metabolism and their combination in iPSC-CM, key processes of cardiac myocytes’ function in health and disease.

## Materials and methods

### Cell culture

#### Cardiac and endothelial differentiation of iPSC

A healthy control iPSC line was used as previously reported [27]. For maintenance, the iPSC were cultured in Essential 8 (E8, Thermo Scientific) and passaged twice a week with Versene solution (Thermo scientific) [28]. For differentiation into CM and endothelial cells (EC), iPSC were passaged onto Geltrex-coated (167 µg/mL, Gibco) 12 Well plates (Sarstedt). Upon reaching 70-85% confluency, cardiac [30] and endothelial [31] differentiation was initiated using a standard published small-molecule differentiation protocol. For iPSC-CM maintenance, the standard RPMI/B27 medium (RPMI # 72400021, B27 #17504044 from Thermo Fisher) and for iPSC-EC, EGM2 medium (#C-22022, Merck) medium was used.

#### 3D-spheroid culture, maintenance and dissociation

3D spheroids were produced as previously described [29]. Shortly, iPSC-CM (day 25 – 28) and iPSC-EC (passage 1 - 3) were dissociated by using 0.25% Trypsin/EDTA (5-7 min) and 0.05% Trypsin/EDTA (1 min), respectively. Cells were counted and transferred to a 15 mL tube containing a ratio of 85% iPSC-CM and 15% iPSC-EC and centrifuged 200 g 5 min. In this study, spheroids consisting of 100.000 cells were produced containing 85.000 iPSC-CM and 15.000 iPSC-EC. The resulting pellet was dissolved in 150 µL RPMI/B27 medium supplemented with 20 % fetal calf serum (FCS, Thermo Fisher), 2 µM Thiazovivin (TZV, Millipore) and 50 ng/mL Vascular Endothelial Growth Factor A (VEGF-A) (#293-VE-010/CF, Bio-Techne) and transferred to a 96-well plate with a conic V-Bottom (#277143, Thermo Fisher). After 24 - 48 h, the medium was replaced with either RPMI/B27 supplemented with 50 ng/mL VEGF-A or MM (see below) supplemented with 50 ng/mL VEGF-A. The medium was refreshed every 2-3 days. To dissolve the 3D-spheroids, 3–5 spheroids were transferred to a 2 mL tube, washed once with 1x phosphate buffered saline (PBS) (#14190-144, Gibco), and incubated with 500 µL 0.25% Trypsin/EDTA for 8-10 min at 37 °C. Subsequently 500 µL FCS was added followed by pipetting until the spheroids macroscopically disintegrate. The dissolved cells were transferred to the appropriate measuring dishes in RPMI/B27 medium supplemented with 20% FCS and 2 µM TZV. After 48 h, medium was replaced with RPMI/B27 or MM. Notably, spheroids were dissolved 8-10 days after spheroid casting. ***Suppl. Table 1 provides the cell numbers and culture formats used for each experiment***.

#### Maturation Medium

For 2D monolayer culture, MM was added at day 25-28. For 3D spheroid culture, the spheroids were cast on day 25-28 and MM was added 1-2 days later. MM was exchanged every three days for 2-3 weeks. The iPSC-CM from these maturation approaches were around day 45 at the time of analysis. For the LT culture, iPSC-CM were kept in RPMI/B27 medium for 70-90 days. For maturation, the MM was composed of two phases. The first 9 days the MM contained FA (palmitate and oleate), GW7647, T3-hormone and dexamethasone. Subsequently, the MM was exchanged to RPMI/B27 supplemented only with FA (palmitate and oleate).

### Molecular analysis

#### Gene expression

Details for RNA isolation, cDNA synthesis, primer sequences and qPCR settings are provided in the supplements.

#### Immunofluorescence

Details are provided in the supplemental methods section.

### AP measurements

IPSC-CM were used for whole-cell patch clamp recordings. The patch-clamp setup consisted of a HEKA electronics system and an EPC10 amplifier (HEKA Elektronik). Microelectrodes with a resistance of 2-4 MΩ were used for the recordings. The microelectrodes were filled with a solution containing (in mmol/L): K-Aspartate 122, KCl 8, NaCl 10, MgCl2 1, HEPES 10, Mg-ATP 5, LiGTP 0.3 (pH 7.2, adjusted with KOH). The bath solution contained in mmol/L (mM): NaCl 140, KCl 4, MgCl2 1, CaCl2 1, Glucose 10, HEPES 5 (pH 7.4, adjusted with NaOH). Action potentials were elicited by current pulses (0.75-1 nA, 2-6 ms) at a frequency of 0.5 Hz at room temperature. Fast capacitance was compensated in cell-attached configuration. Recordings started after cell stabilization, which was approximately 5 min after rupture. The access resistance was typically <10 MΩ after rupture. The signals were filtered with 2.9 and 10 kHz Bessel filters and recorded using the EPC10 amplifier. The recordings were then analyzed using LabChart8.

### Live cell imaging

#### Cytosolic Ca^2+^ measurements

Ca^2+^ measurements were performed on single cells in 2D (spheroids were dissociated as previously described). The cells were loaded with Indo-1 (stock 1µg Indo-1 in 1 µL Pluronic-F127) in Tyrode solution containing in mM: 140 NaCl, 5.4 KCl, 1 MgCl_2_, 1.8 CaCl_2_, 10 Na-HEPES, 10 glucose; pH = 7.4). Microscopy set-up consisted of a customized manual and automatic IonOptix set-up (Milton, USA) coupled with an inverted epifluorescence microscope (Eclipse Ti, Nikon). The experimental set-up followed the sequential steps: 1) 0.25 Hz stimulation 2) 0.5 Hz stimulation 3) 1.0 Hz stimulation 4) 0.5 Hz stimulation, allowing cells to acquire pace, 5) addition of 50 nM Isoproterenol (Iso) solution (at 0.5 Hz pacing) 6) exchange for 100 nM Iso solution (at 0.5 Hz pacing). The IonOptix set-up allows one to measure the same set of cells sequentially, therefore individual cell responses can be normalized to the cell’s own baseline activity, useful for applying paired statistical comparisons.

#### Mitochondrial Ca^2+^ imaging

The iPSC-CM were transfected using Lipofectamine LTX Kit (#<u>A12621</u>, Thermo Fisher) mixing 300 µL OptiMEM (#51985034, Thermo Fisher) with 1.5–2 µg mitoPericam-plasmid [30], 12 µL lipofectamine LTX reagent and 1µL PLUS reagent (5 min incubation), which was subsequently added to the cells cultured in 1.7 mL RPMI/B27 medium. After 6 h, the medium was refreshed and after 2 days the cells were imaged using aconfocal microscope (Leica SP8) with the following settings: 63x water objective, sequential scan, excitation with 405 nm (8%) and 488 nm (14 %) single channel detection using photo-multiplier tube at 500-550 nm, gain 900, offset -1, zoom 2 and resolution of 1352x1352. The cells were measured in Tyrode solution (containing in mM: 140 NaCl, 5.4 KCl, 1 MgCl_2_, 1.8 CaCl_2_, 10 Na-HEPES, 10 glucose; pH = 7.4) at room temperature with 0.5 Hz pacing.

#### Mitochondrial membrane potential

The iPSC-CM were loaded with 2.5 nM Tetramethylrhodamin, Methylester, Perchlorat (TMRM) (Thermo Fisher) and 100 nM MitotrackerGreen (Thermo Fisher) (in RPMI/B27) and incubated for 1 h at 37 °C in the dark. Afterwards, the medium was exchanged with RPMI/B27 with 2.5 nM TMRM and imaged with a confocal microscope (Leica SP5) at 37 °C. Whole images (63x oil objective) were analyzed for the channels of TMRM and MitotrackerGreen for their mean pixel intensity in Image J, and a ratio of TMRM/Mitotracker was calculated.

### Transmission electron microscopy

The 3D spheroids were washed once with 1x PBS and fixed with Karnovsky’s fixative overnight at 4 °C. The next day, the fixative was replaced with cacodylate buffer (0.1 M, pH 7.4). Samples were stained using 1% osmium tetroxide and 1% uranyl acetate, dehydrated using graded concentrations of ethanol, embedded in Durcupan resin using a laboratory microwave, and sectioned. Thin sections (70 nm) were mounted on copper grids, post-stained with 3% lead citrate, and imaged using a Talos L120C transmission electron microscope (Thermo Scientific) operated at 120kV, equipped with a 4K × 4K Ceta CMOS Camera (Thermo Scientific).

### Statistics

For comparisons of two groups, the Mann–Whitney *U* test was performed. For multiple comparisons analysis of variance (ANOVA) or Kruskal-Wallis test, corrected with the Dunnett’s method was applied. For testing two groups with more than one value per cell, a two-way ANOVA with Geisser-Greenhouse’s correction was used. *P* values are two-sided and considered statistically significant if *p* < 0.05 (* p<0.05, ** p<0.01, ***p<0.001). Analysis was performed with GraphPad Prism 9. Details for each statistical test are provided in the figure legends.

## Results

### A combinatorial 3D-spheroid in MM approach to boost iPSC-CM maturation

A control iPSC line from a healthy donor was differentiated using a standard small-molecule Wnt-activator/Wnt-inhibitor protocol to produce ventricular iPSC-CM [31]. At days 25-28, the cultures were randomized into different maturation conditions (Figure 1A, B). RPMI/B27 served as a standard condition medium and long-term (LT) culture in RPMI/B27 for 60-90 days was used as a reference for an established maturation strategy providing cells with a well-defined phenotype. The quality of the cardiac differentiations was verified by a high proportion of CM with 91% ±2% (SEM) cTNT positive cells (Suppl. Fig. 1A). To further demonstrate the specific differentiation into ventricular CM, we demonstrated a predominant abundance of the ventricular marker myosin light chain ventricle (*MYL2)* concomitant with low expression of the atrial marker *NR2F2* (Suppl. Fig. 1B). Further, sub-type-specific analysis using immunofluorescence confirmed that iPSC-CM are positive for the ventricular marker MLC2v and negative for the atrial marker myosin light chain atrium (MLC2a; Suppl. Fig. 1C). For forming 3D, organoid-like spheroids, CM and EC were cast in a proportion of 85% CM to 15% EC, as previously described [29]. Whole-mount stainings of the 3D-spheroids stained against muscle (alpha-actinin) and EC (CD31) marker proteins confirmed the presence of both cell types 15 days after spheroid casting, with relative volume occupancy indicating that EC may be able to proliferate within spheroids. Furthermore, we observed regularly structured z-discs in CM and EC membrane protrusions linking to adjacent CM (Fig. 1C). Notably, the spheroids were dissolved for subsequent analyses on the single-cell level. At this point, the iPSC-EC were vastly lost and did not survive the rigorous trypsinization process.

**Fig. 1:**
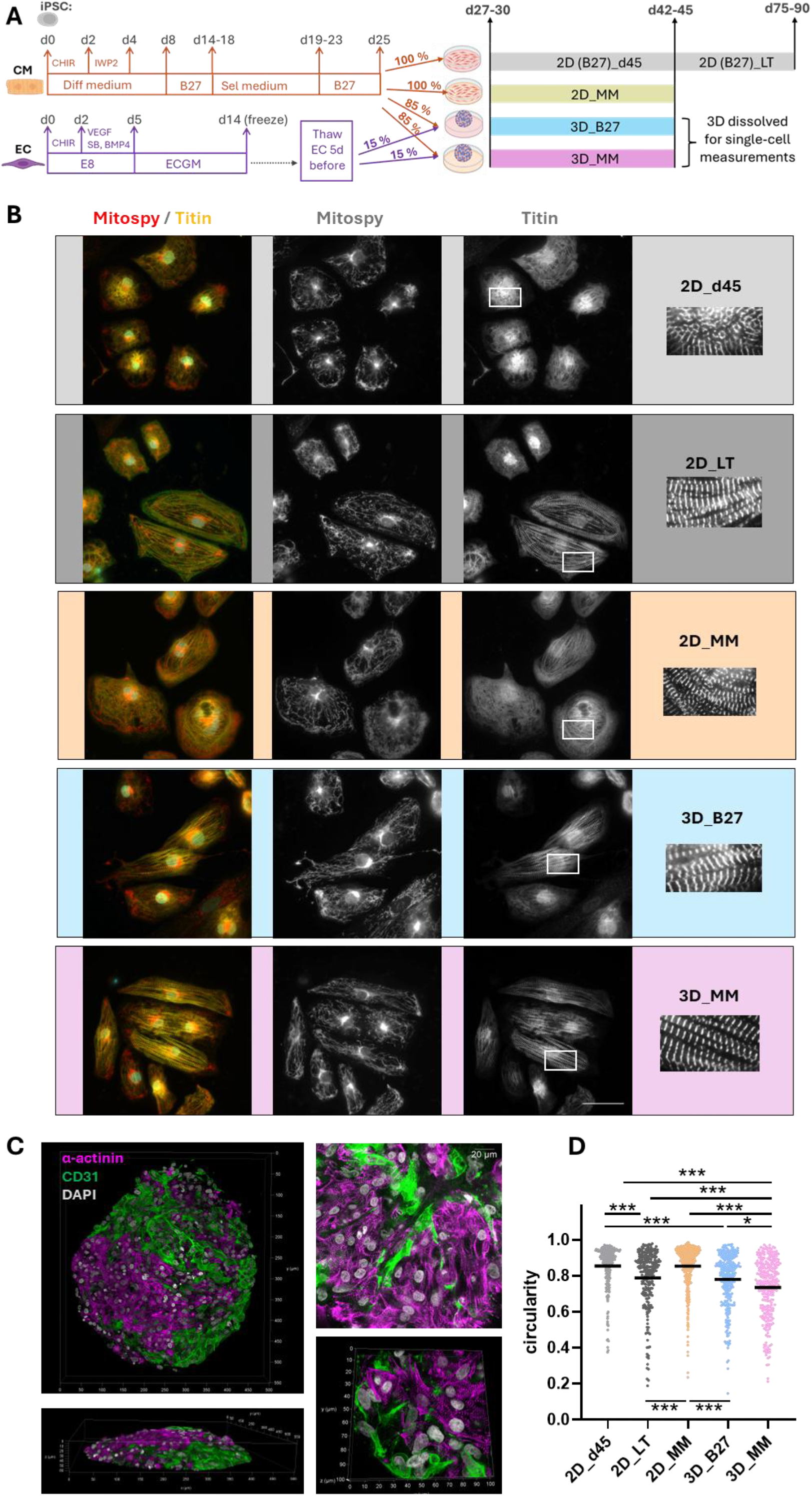
Experimental design and structural and mitochondrial investigation. **A:** Schematic summary of maturation conditions used. **B:** Exemplary staining to visualize the mitochondrial network with Mitospy (red) and the sarcomere with titin-M line (yellow) and DAPI counterstaining to visualize the nucleus (cyan). For visualization purposes, the brightness and contrast has been enhanced. **C:** Confocal microscopy of whole spheroids stained with antibodies against α-actinin (purple) and CD31 (green), and using the nuclei stain Hoechst (grey). The images on the left-hand side provide volumetric images of the spheroid surface (40x water immersion objective). The images on the right depict representative CM and EC within the spheroid at high resolution (63x glycerol objective). **D:** Quantification of cell shape with circularity index. No. of differentiations /analyzed cells for 2D_d45 [4/304], 2D_LT [3/250], 2D_MM [4/395], 3D_B27 [4/280] and 3D_MM [4/283]. P-values were calculated by Kruskal-Wallis test with Dunn’s multiple comparisons against every group. Data is represented as beeswarm plots with mean. Significant p-values are marked by * p<0.05, ** p<0.01, ***p<0.001.

The novel-composite maturation medium (MM) contains the standard B27 supplement (in RPMI, Glutamax, 2g/L glucose) together with the hormones T3 and dexamethasone, a PPARα-agonist and two BSA-complexed fatty-acids (FA), palmitate and oleate. In the adult heart, energy is mainly provided from the oxidation of FA in mitochondria [32]. Since immature iPSC-CM, as fetal or neonatal CM, rely primarily on glycolysis and glucose oxidation, metabolic maturation toward an adult phenotype requires a transition from glucose to FA oxidation [3]. In fact, in a 2D culture, the MM increased gene expression of *PDK4* and *CPT1B* (Suppl. Fig. 1D), indicative of a shift from glucose to FAO. In contrast, MM alone neither affected expression of the Na^+^ channel *SCN5a* or the K^+^ channel *KCNJ2* (Suppl. Fig. 1E), nor did it induce a switch of sarcomeric genes from an embryonic to an adult isoform, i.e., myosin heavy chain (MHC) from *MYH6* to *MYH7*, or titin from the longer titin-*N2BA* to the *N2B* isoform (Suppl. Fig. 1F). Concomitant with the increased expression of metabolic genes, MM increased the mitochondrial membrane potential gradient (ΔΨ_m_; Suppl. Fig. 1G) as the driving force for ATP production via the F_1_F_o_-ATP synthase. These data suggest that MM alone promotes metabolic maturation, but not EC coupling, in iPSC-CM.

To further improve maturation, we combined MM with 3D-spheroid culture, in which 3D-spheroids are cast at d25-28, MM treatment is initiated on d27-30 and continued for 14 days. In the following, we assessed whether the application of MM to 3D-spheroids drives a superior maturation beyond the benefits of MM or 3D-spheroids alone, respectively, and compared to the well-defined benefits of LT culture. To systematically assess this, the following maturation conditions were analyzed: (1) 2D culture with MM, (2) 3D-spheroid culture in standard RPMI/B27 medium, and (3) 3D-spheroid culture with MM. To compare the maturation progress, the following two conditions were used as reference states: (4) 2D culture in RPMI/B27 medium, which was compared to the d45 timepoint of all maturation approaches, and (5) LT culture in RPMI/B27 medium for 60-90 days, serving as reference for the matured iPSC-CM stage.

### CM within 3D-spheroids cultured in MM show a mature cell shape

We first analyzed the cellular structure by visualizing the sarcomeric and mitochondrial structure and number of nuclei (Fig. 1B). Mature human CM are characteristically multi-nucleated rod-shaped cells with unidirectional, aligned sarcomeres. Here, iPSC-CM cultured in B27 appeared as round cells with regular sarcomeres that follow the oval shape of the cell, which is the typical presentation of an iPSC-CM. The cell shape shifted toward the adult rod shape when the 3D maturation cue was introduced. Accordingly, the circulation index (a value of 1 indicates a perfect circle) significantly decreased with enhanced maturity with the lowest value for 3D_MM, followed by 3D_B27 and LT, while 2D_B27 and 2D_MM show the highest, i.e., less mature values (Fig. 1D). In an unloaded condition, the sarcomere length of an adult human CM is 1.78 µm [33]. By the LT conditions, this length is nearly achieved with a mean sarcomere length of 1,74 µm, followed by 3D_MM (mean 1.68 µm) and lower values for 2D_MM (mean 1.65 µm) and 3D_B27 (1.66 µm) (Suppl. Fig. 2A). Furthermore, the percentage of multi-nucleated cells was not different between the tested conditions (Suppl. Fig.2B). Next, we quantified the mitochondrial network using Mitospy for mitochondrial staining (Fig. 1B). We did not observe any impact of spheroid culture or MM on the mitochondrial network structure, which was a diffuse meshwork with 13-14% of connected network structures (Suppl. Fig. 2C). In all conditions, mitochondria did not align with the sarcomere (Fig. 1B).

### Emerging ultrastructural features in matured iPSC-CM

Using electron microscopy (EM) of whole cardiac spheroids (3D_MM), advanced structural organization characteristic of mature CM could be observed, including well-defined sarcomeric Z-discs (green arrows) and aligned myofibrils (Fig. 2). Evidence of mechanical coupling between CM via desmosomes and adherens junctions was observed, suggesting the development of structural integration of cells within the spheroid (blue arrows). Additionally, we observed numerous mitochondria with well-defined cristae, reflecting metabolic maturation (orange arrows). Interestingly, several invaginations of the plasma membrane, structurally compatible with t-tubules, were observed in 3D_MM CM (pink arrows), although these were not as frequent as in adult CM. Overall, 3D_MM conditions promoted ultrastructural maturation, supporting improved metabolism and EC coupling.

**Fig. 2:**
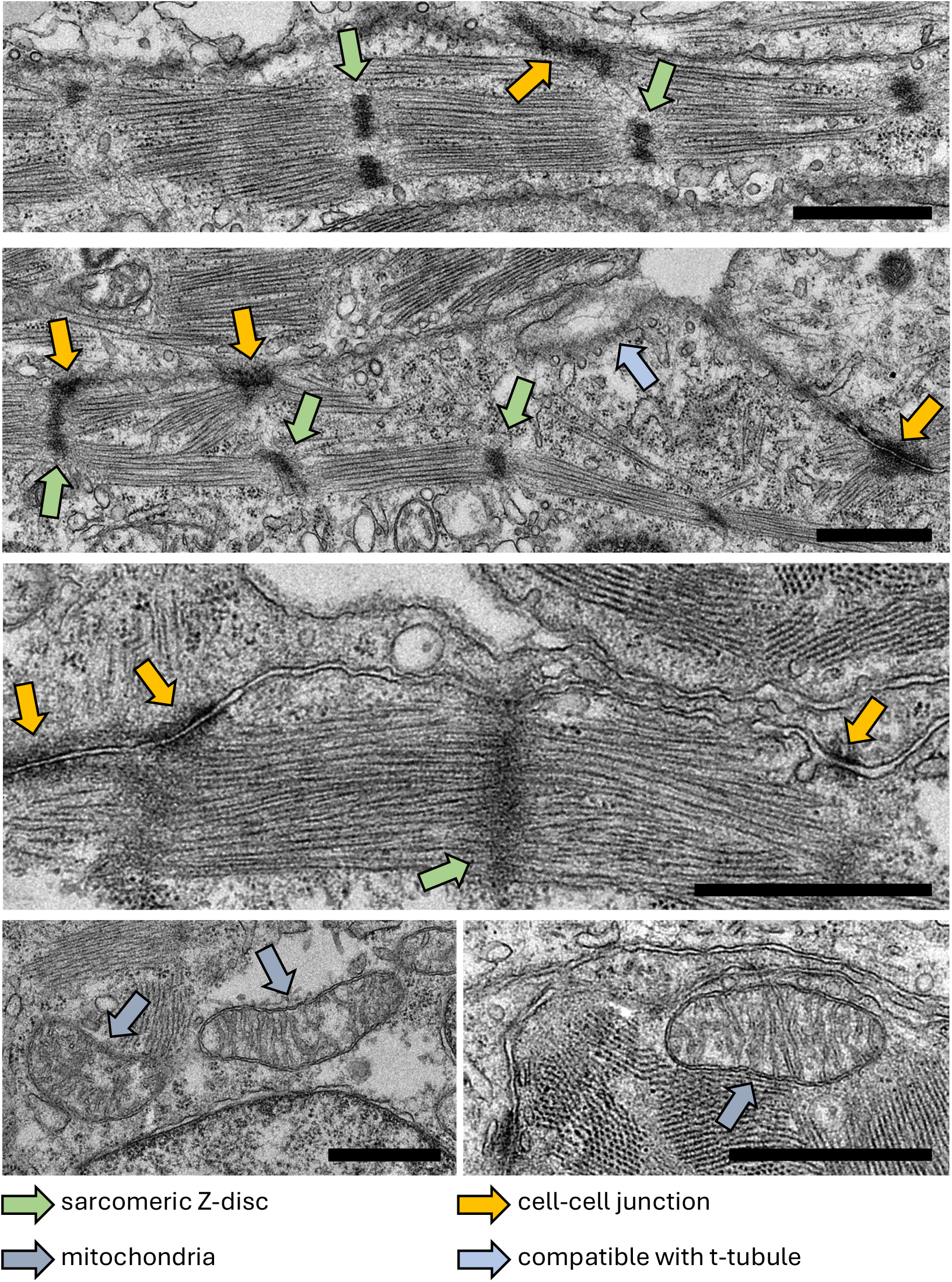
Ultrastructural analysis of 3D_MM spheroids. Representative transmission electron micrographs revealed aligned and well-organized myofibrils in whole 3D_MM spheroids (green arrows indicate Z-discs), evidence of mechanical coupling between CM (desmosomes and adherens junctions, blue arrows). Several invaginations of the plasma membrane reminiscent of developing transverse (t-) tubules were also observed (pink arrow). Orange arrows highlight mitochondria with well-defined cristae. Scale bars = 1 µm.

### Electrophysiological properties of CM obtained from 3D_MM are comparable to those of CM obtained from LT culture

Next, we assessed the electrophysiological properties of iPSC-CM matured under different conditions (Fig. 3A). Adult human ventricular CM have a RMP between -80 mV and -90 mV, an APD of 300-450 ms and a characteristic AP shape with a rapid upstroke, a prominent plateau phase followed by repolarization resulting in a peak-and-dome shape [24, 34]. Single-cell patch-clamp recordings revealed a more negative RMP for 3D_MM (mean -71.3 mV) compared to 2D_MM (mean -67.6 mV) and 3D_B27 (mean -67.9 mV), accompanied by the highest AP amplitude (APA) for 3D_MM (mean 123.3 mV) compared to 2D_MM (mean 116.0 mV) and 3D_B27 (mean 117.2 mV) (Fig. 3B). However, only CM from 2D_MM reached the adult-like APD range of 300 - 450 ms, and their AP resembled most closely the AP shape of adult ventricular human cardiomyocytes (Fig. 2A/B). It should be noted that the APD significantly differed between the different maturation conditions, with 3D_MM exhibiting a longer APD90 (mean 580 ms) than 2D_MM (mean 415 ms), while 3D-iPSC-CM without MM showing the longest APD90 (mean 635 ms; Fig. 3B). The maximal upstroke velocity was highest for 3D_MM (mean 101.7 mV/ms), followed by 2D_MM (mean 77.1 mV/ms) and 3D_B27 (66.2 mV/ms). All data are summarized in Fig. 3C. LT culture also drives CM maturation towards electrophysiological properties closely resembling those of adult CM, as previously demonstrated [18]. Specifically, iPSC-CM matured by LT culture showed AP with a typical peak-to-dome shape, an APD90 of 260 ms, and an RMP of -75 mV [18].

**Fig. 3:**
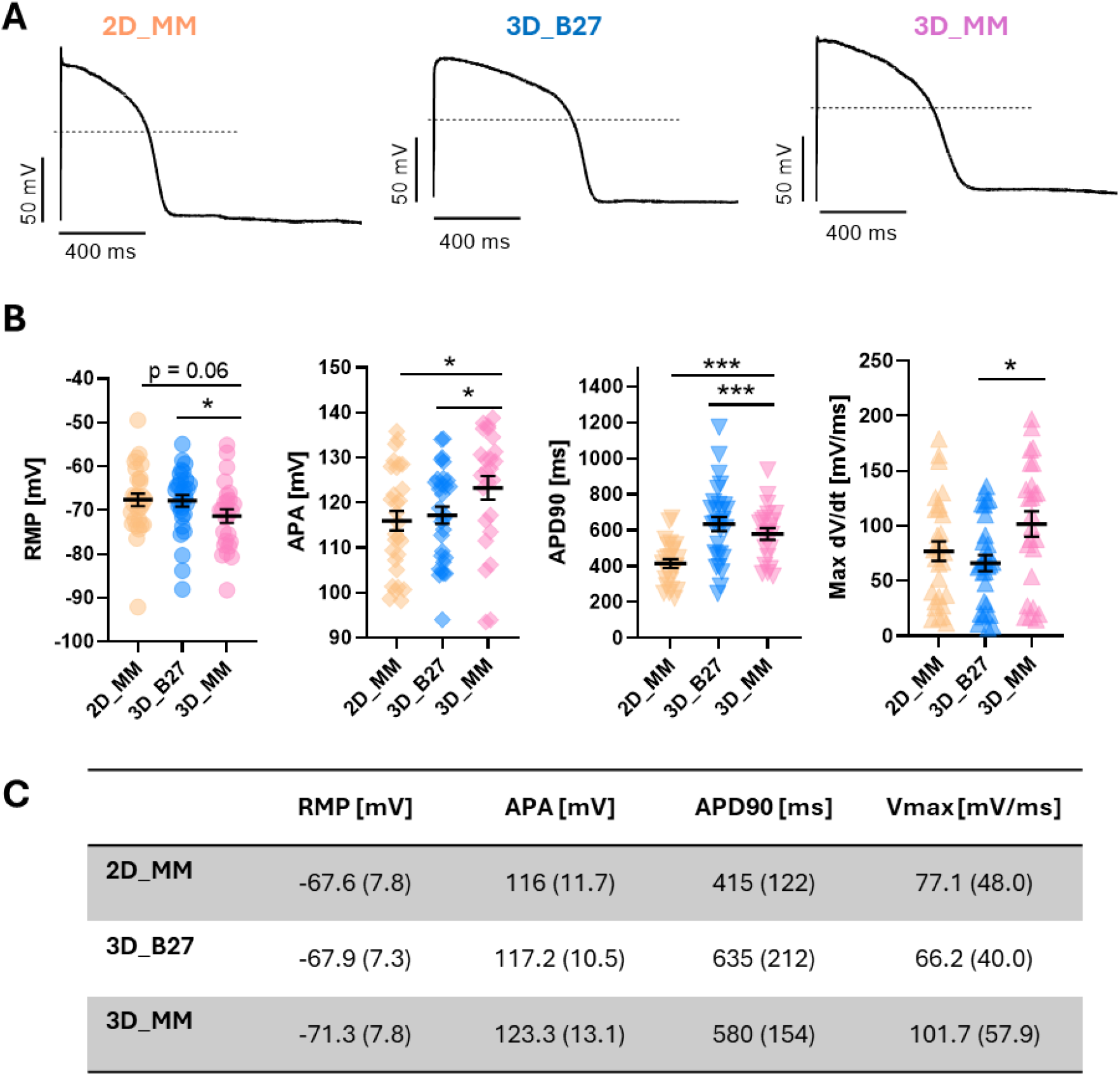
Electrophysiological investigation of the different maturation protocols. **A:** AP recordings for 2D_MM, 3D_B27 and 3D_MM at 0.5 Hz stimulation. **B:** Analysis of the different AP parameters at 0.5 Hz: RMP (resting membrane potential), APA (action potential amplitude), APD90 (action potential duration for 90% of the AP) and the maximum upstroke velocity (max dV/dt). [Number of differentiations/analyzed cells] for 2D_MM [3/29], 3D_B27 [3/30] and 3D_MM [3/25]. Statistics by Mann-Whitney test for 3D_MM vs 2D_MM and 3D_MM vs 3D_B27. Significant values are marked * p<0.05, ** p<0.01, ***p<0.001. **C:** Summary of data for the different AP parameters. Shown is: mean (Stdev).

Collectively, these data show that the combination of 3D_MM provides the most consistent improvement in cardiac AP features, with 3D_B27 alone yielding cells with electrophysiological properties indicative of a lower degree of the maturation. The maturation state after 3D_MM is comparable to the effects of LT culture, but is achieved earlier.

### Ca^2+^ homeostasis and drug sensitivity are improved by the combinatorial maturation approach

Ca^2+^ is the key ion in EC coupling that activates the myofilaments for contraction. Two major mechanisms to increase cardiac contractility are β-adrenergic stimulation and frequency-dependent force potentiation, also known as the “Bowditch effect” or “force-frequency relation” [35]. Under either condition, the increase in force is driven by an increase in the amplitude of cytosolic Ca^2+^ transients [35]. Therefore, we determined cytosolic Ca^2+^ concentrations ([Ca^2+^]_c_) with the ratiometric Ca^2+^ dye Indo-1 and either increased the stimulation frequency (0.25 Hz to 1.0 Hz) or stimulated β-adrenergic receptors with two subsequent Iso concentrations (50 nM and 100 nM). Diastolic [Ca^2+^]_c_ was similar in CM of 2D_MM and all 3D conditions and overall lower than in 2D_LT (Fig. 4A). In contrast, systolic [Ca^2+^]_c_ and the amplitude of [Ca^2+^]_c_ transients were increased in CM of 3D_MM compared to 2D_MM and 3D_B27 (Fig. 4B, C). The frequency-dependent potentiation of systolic [Ca^2+^]_c_ was increased by MM in 2D or 3D compared to the non-MM conditions (3D_B27 and 2D_LT; Fig. 4D, Suppl. Fig. 3A, B). Regarding β-adrenergic responsiveness, the 3D_MM condition showed the strongest increase in systolic [Ca^2+^]_c_ and the fastest decay of the [Ca^2+^]_c_, followed by the 3D_B27 and 2D_MM conditions (Fig. 4E, F; Suppl. Fig. 3B-F). An overview of all values is provided in Fig. 4G.

**Fig. 4:**
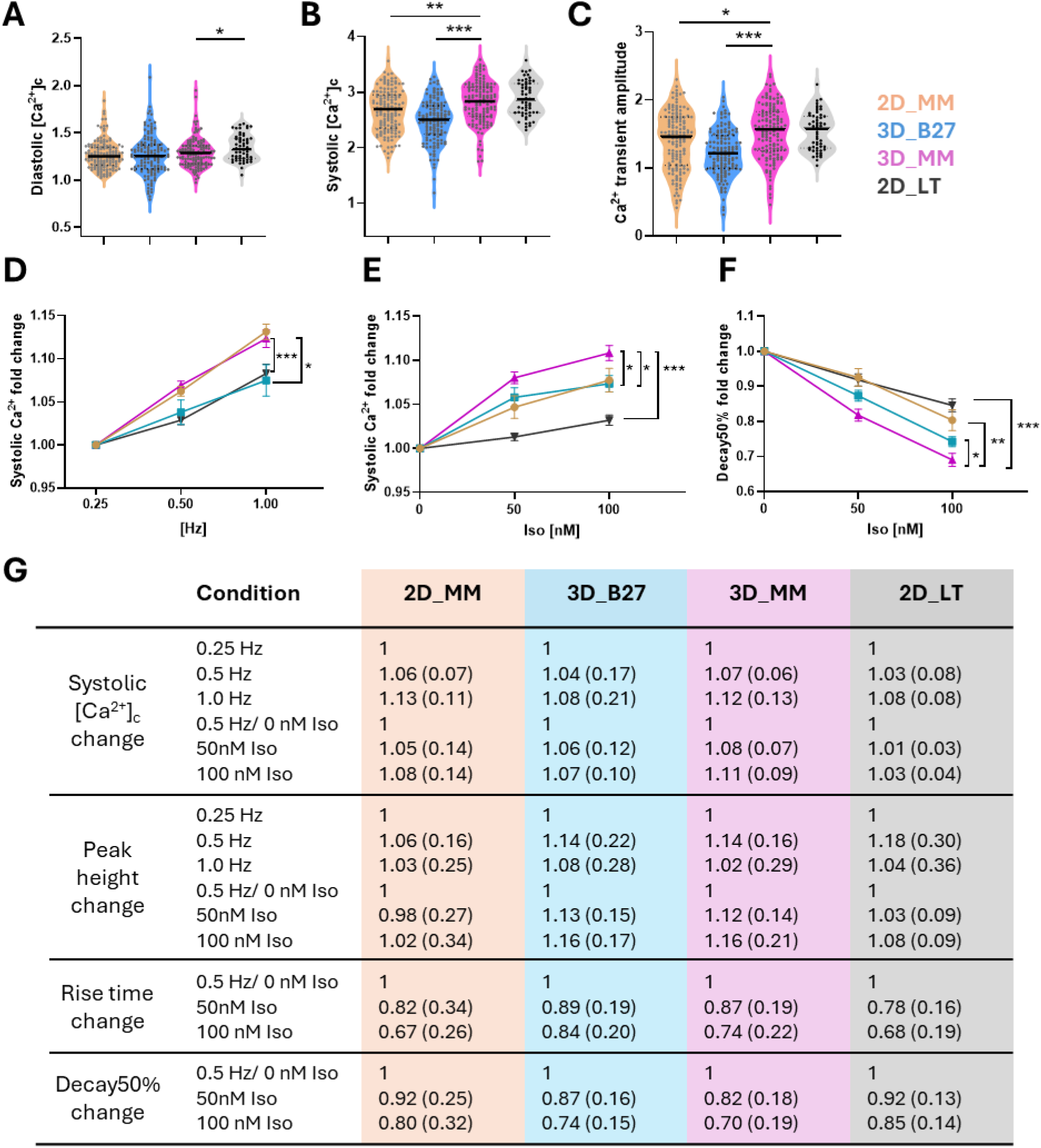
3D_MM improved Ca^2+^ homeostasis and sensitivity to electrical and β-adrenergic stimulation. Measurement of cytosolic Ca^2+^ concentrations ([Ca^2+^]_c_) with the ratiometric Ca^2+^ indicator Indo-1 AM (data as ratio 405/495). **A-C:** Analysis of diastolic [Ca^2+^]_c_ **(A)**, systolic [Ca^2+^]_c_ **(B)**, [Ca^2+^]_c_ amplitude height **(C)** at 0.5 Hz pacing for different maturation conditions. P-value by Kruskal-Wallis test with Dunnett’s multiple comparisons against 3D_MM. **D:** The iPSC-CM from different maturation conditions were subjected to increasing pacing (0.25 Hz to 0.5 Hz to 1Hz). Since matching cells were measured, the basal condition is set to 1 and data is shown as fold change. Statistics was performed by Two-way ANOVA with Geisser-Greenhouse’s correction. P-value (column factor) for 3D_MM vs 2D_MM; 3D_MM vs 3D_B27 and 3D_MM vs 2D_LT. **E, F:** The iPSC-CM from different maturation conditions were subjected to increasing Iso concentrations (0 nM to 50 nM to 100 nM) with 0.5 Hz pacing. Since matching cells were measured, the basal condition is set to 1 and data is shown as fold change of the respective Ca^2+^ parameter. Statistics was performed by Two-way ANOVA with Geisser-Greenhouse’s correction. P-value (column factor) for 3D_MM vs 2D_MM; 3D_MM vs 3D_B27 and 3D_MM vs 2D_LT. **A-F:** [No. of differentiations /analyzed cells] for 2D_MM [5/145], 3D_B27 [5/137], 3D_MM [5/137] and 2D_LT [2/57]. Significant values are marked * p<0.05, ** p<0.01, ***p<0.001. **G**: Summary of all data points assessed with different pacing and Iso-treated conditions. Data is shown as: mean (StDev). Iso = Isoprenaline.

Taken together, especially the combined 3D_MM protocol significantly advances maturation in respect to cytosolic Ca^2+^ handling, substantially boosting frequency-dependent potentiation of [Ca^2+^]_c_ transients and the β-adrenergic response.

### Reliable measurements of mitochondrial Ca^2+^ content in matured iPSC-CM using a genetically encoded mitochondrial Ca^2+^ indicator

To match energy supply with demand during transitions in workload, Ca^2+^ is taken up into mitochondria via the mitochondrial Ca^2+^ uniporter (MCU) to stimulate Krebs cycle dehydrogenases, which accelerates the regeneration of NADH and FADH_2_ as electron donors for the respiratory chain [2, 36]. For an efficient transfer of Ca^2+^ from the SR (the main source of systolic cytosolic Ca^2+^) to mitochondria, a close localization of mitochondria to the SR and the formation of so-called “Ca^2+^ microdomains” is required [36]. An important driving force for mitochondrial Ca^2+^ uptake is the mitochondrial membrane potential (ΔΨ_m_), which we determined with tetramethylrhodamine (TMRM). Metabolic maturation protocols in 2D or 3D increased ΔΨ_m_ compared to non-MM conditions (3D_B27; Fig. 5A). To monitor mitochondrial Ca^2+^ concentrations, we transfected iPSC-CM with plasmids encoding mitoPericam [37], a mitochondria-targeted Ca^2+^ reporter (Figure 5B). Under steady-state conditions (0.5 Hz pacing), mitochondrial Ca^2+^ concentrations were slightly higher in 3D_MM compared to 2D_MM conditions (Fig. 5C). Since there was no difference in ΔΨ_m_ between these conditions (Fig. 5A), the higher mitochondrial Ca^2+^ concentrations in 3D_MM might be related to higher cytosolic Ca^2+^ transient amplitudes (Fig. 4B) and/or a more mature spatial organization of the elements involved in EC coupling (Fig. 1B).

**Fig. 5:**
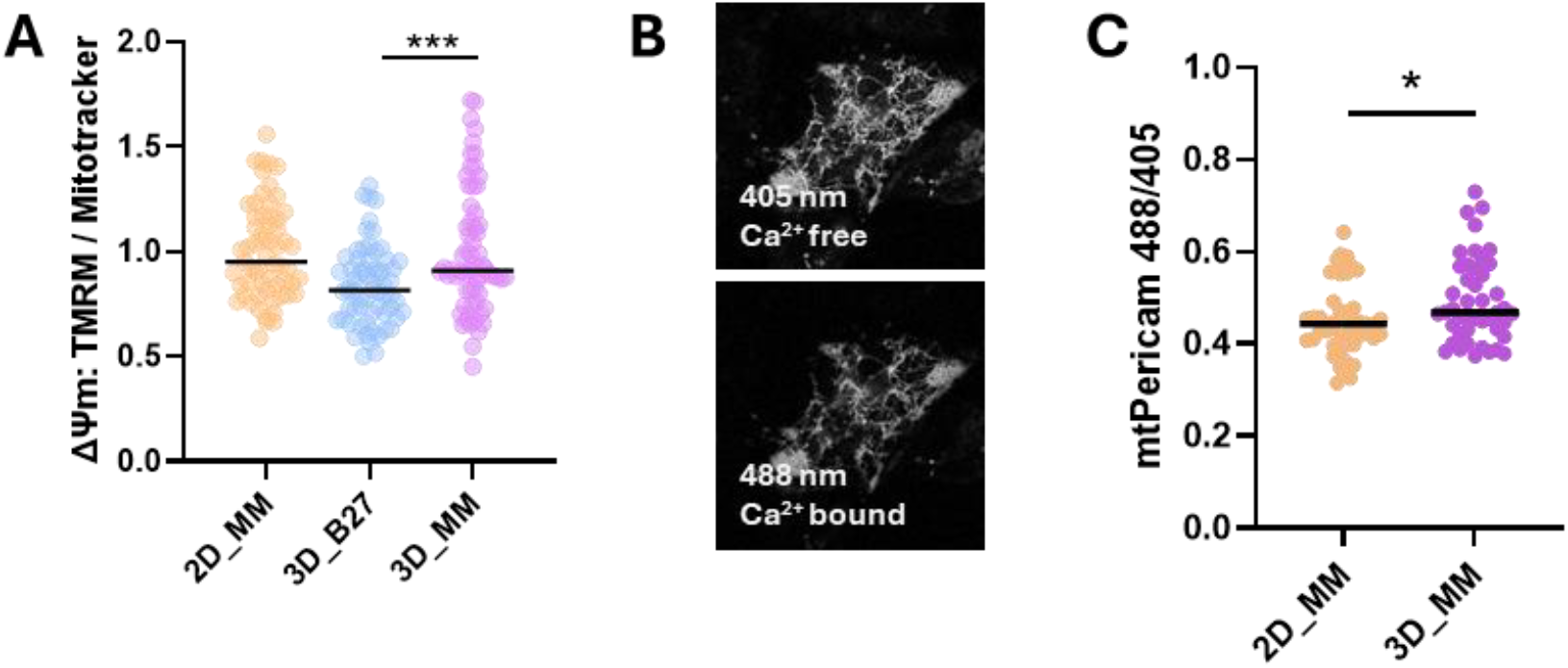
Mitochondrial Ca^2+^ analysis in matured iPSC-CM. **A:** Measurement of mitochondrial membrane potential ΔΨ_m_. [Cardiac differentiations /No. of pictures analyzed] for 2D_MM [3/61], 3D_B27 [3/64] and 3D_MM [3/65]. P-values by ordinary one-way ANOVA with Dunnett’s multiple comparisons against 3D_MM. Significant p-values are marked by * p<0.05, ** p<0.01, ***p<0.001. **B:** Representative images of iPSC-CM transfected with the mitoPericam plasmid. Excitation with 405 nm visualizes unbound Pericam and 488 nm the Ca^2+^ bound Pericam. **C:** Quantification of the Pericam signal as a relative measure of mitochondrial Ca^2+^. [Cardiac differentiations /No. of pictures analyzed] for 2D_MM [3/49] and 3D_MM [3/45]. P-value by Mann-Whitney test. Significant p-values are marked by * p<0.05, ** p<0.01, ***p<0.001.

## Discussion

Production of mature adult iPSC-CM remains a complex challenge; however, it is required for optimized disease modelling and cardiotoxicity testing. Here, we report a combinatorial approach for culturing iPSC-CM as 3D spheroids in a defined MM to significantly enhance CM properties, which does not require specialized equipment. 3D_MM enhanced the structural, electric and Ca^2+^ handling physiology of the iPSC-CM with the strongest effect on cell shape, APA, RMP, systolic [Ca^2+^]_c_ and β-adrenergic responsiveness. It had weaker effects on ΔΨ_m_, with no effect observed on sarcomeric length or the mitochondrial network, compared to control conditions. Notably, in some aspects, the single approaches were as successful as the combination approach 3D_MM, for example, when considering changes in ΔΨ_m_; therefore, the use of single maturation cues could be considered for specific applications. However, in our study, only the 3D_MM approach provided comprehensive structural, functional and metabolic maturation. This underscores the emerging principle of integrating multiple strategies instead of exploring the “form follows function” principle, i.e., that pushing one aspect of maturation will lead to overall maturation). In line with previous publications, LT culture produces iPSC-CM with mature cell size, shape and sarcomeric arrangements [16, 38]. Compared to LT, the 3D_MM condition achieves similar maturation benefits in structure and AP physiology, and even outperformed LT in respect to achieving a mature cell shape and Ca^2+^ handling properties.

The cell shape shifts towards a rod shape with 3D_MM, with improvements also seen in 3D_B27. To date, the CM morphologies most closely matching adult cells *in situ* have been achieved with nanofibers [39] and nanopatterns [13, 40]. EM images of iPSC-CM on nanofibers show well-aligned myofilaments that are comparable to our 3D_MM condition [39]. An advantage of the 3D_MM, however, is that more iPSC-CM can be produced for assays that require a higher cell number, such as for mRNA or protein analyses.

Further, our 3D_MM protocol produces CM with RMP and AP parameters comparable to CM from LT [18], whereas maturation approaches that use electrical pacing for maturation yield cells with adult-like AP [13, 23]. In future studies, electrical stimulation could be integrated into 3D_MM as an additional cue to push maturation. It remains a major challenge to optimally combine multiple maturation cues to achieve synergistic maturation boosts. This is complicated by the possibility of different dosages of the individual cues, as well as their timing (chronic vs transient) and order (simultaneously or sequentially). For example, combining our 3D-spheroids with a 1 Hz stimulation period resulted in cell damage (data not shown). Other groups reported transient pacing periods starting at 2 Hz in young iPSC-CM, followed by 1 Hz [13]. Unravelling these details further will hopefully produce iPSC-CM that come even closer to the human adult CM.

While human CM have defined absolute values for RMP and APD, Ca^2+^ parameters are often measured and reported as relative values, which makes them harder to be compared between different experimental groups. In general, immature iPSC-CM have lower systolic [Ca^2+^]_c_ levels and are less sensitive to drug treatments due to lower ion channel densities and because they are lacking ultrastructural features like t-tubules and intercalated discs, which functionally link ion channels, the SR and mitochondria [41, 42]. As the 3D_MM protocol matures CM in respect to cell shape and AP, there is also concomitant refinement in Ca^2+^handling (higher systolic [Ca^2+^]_c_ levels). Apart from basal Ca^2+^ parameters, correct and sensitive responsiveness to electrical stimuli and known drugs is better defined. To date, drug tests and screenings primarily focus on multi-electrode assays with beating rate and field potential duration as readouts, enabling high-throughput screening [43]. Here, we report a comprehensive analysis of Ca^2+^ parameters in matched cells with sequentially increasing pacing rates and Iso stimulation. Low Iso doses of 10 nM are reported to increase the beating rate of iPSC-CM [44, 45], whereas higher Iso concentrations up to 1 µM are necessary to provoke an observable Ca^2+^ response [31, 46]. In general, an advanced maturation state of iPSC-CM correlates strongly with their ability to capture drug responses sensitively and correctly [13, 42]. In our study, the underlying Ca^2+^ cycling in matured iPSC-CM reacts to increased pacing and Iso treatment in a dose-dependent manner. 3D_MM respond to 50 nM Iso with higher systolic [Ca^2+^]_c_ and faster Ca^2+^ cycling compared to all other conditions, and even larger effects were achieved upon treatment with 100 nM Iso. The observed sensitivity to Iso indicates the presence of mature microdomains, since an adequate inotropic Iso response has been described to require a functional dyad organization [42, 47]. Since 3D_MM can yield high cell numbers without specialized equipment, it represents a suitable approach to produce matured iPSC-CM for use in drug screenings and toxicity tests.

3D_MM benefits iPSC-CM on many physiological aspects; however, the least obvious effects were observed when analyzing mitochondrial architecture and function. For an effective mechano-energetic coupling, the mitochondria align with the sarcomere in adult CM to supply sufficient ATP to the high energy demand of contracting CM, and are also in close contact with the SR to allow efficient SR-mitochondrial Ca^2+^ transmission [2, 48]. To date, this mitochondrial alignment has not yet been achieved, and iPSC-CM maturation can be enhanced with respect to oxidative capacity, FAO and ΔΨ_m_, but the mitochondrial architecture remains a diffuse network of mitochondria, and this aspect was not resolved when applying our 3D_MM protocol. Since iPSC-CM respond to stimuli that promote CM maturation *in vivo* (e.g., hormone stimulation, 3D culture conditions), a deeper understanding of mitochondrial biogenesis during cardiac embryogenesis might help identify an effective cue.

### Limitations

A major limitation is the subpar dissociation of the spheroids into 2D format. A 7-10 min incubation step with 0.25% Trypsin/EDTA was able to sever the spheroid into single cells and cell clusters. However, this was accompanied by significant cell loss, whereas milder reagents such as Versene or Accutase were not strong enough to separate the cells. Furthermore, iPSC-CM slowly change their rod-like cell shape back to a more rounded state once they are removed from the spheroid (over the course of days). This may mask or dilute the maturation benefits of spheroids when cells are measured in 2D. Therefore, the unexpectedly poorer performance of 3D_B27 in AP and Ca^2+^ handling compared to 2D_MM could be a result of the difficulty in dissociating cells from spheroids. This would also suggest that 3D_MM performs even better if a robust dissociation protocol is developed that improves cell survival and preserves maturation state more effectively.

## Author contributions

SR developed the MM composition, generated the 3D spheroids, performed most of the experiments, drafted and wrote the manuscript, and received funding for this study. HE performed the Ca^2+^ measurements. NW and JU generated the qPCR, sarcomeric and mitochondrial data. ND performed the AP measurements. FSW and ERZ provided the whole-spheroid and TEM images. KSB developed the concept, received funding, and wrote the manuscript. CM contributed to editing the MS.

## Acknowlegdements

We acknowledge the SCI-MED imaging facility at the Institute for Experimental Cardiovascular Medicine in Freiburg for providing access to the confocal microscope, which was funded by the German Research Foundation (DFG - Projektnummer 404198760). We thank Dr. Josef Madl for his support with imaging, and Jonas Heer for help with immunohistochemistry.

