## Supplemental Information for "Enhanced iPSC Cardiomyocyte Maturation via Combined 3D-Culture and Metabolic Cues"

### Supplemental Information Rebs et al.

#### Supplemental Methods:

For analysis of structural, electrophysiological and $Ca^{2+}$ handling parameters the cells were digested and reseeded according the specific requirements of the read-outs. Digestion of 2D-monolayer and dissociation of 3D-spheroids is described in the method section: *3D-spheroid culture, maintenance and dissociation*. Suppl. Tab. 1 summarizes the cell and spheroid numbers used for the individual assays performed in this study.

***Suppl.Tab. 1: Summary of cell and/or spheroid number used for specific assays.***

| Assay | Format | 2D cells counted | No of spheroids |
|---|---|---|---|
| IF | 20 mm coverslip in 12 well plate | 30.000 – 40.000 | 2 - 3 |
| EM | whole spheroids | 100.000 | 3 - 4 |
| AP | 35 mm dish | 100.000 | 3 - 4 |
| $Ca^{2+}$ | 35 mm dish | 150.000 | 3 - 5 |
| Mito-$Ca^{2+}$ | 35 mm dish | 300.000 | 6 - 8 |
| Coating: 167 µg/mL Geltrex for a minimum of 30 min at 37 °C | | | |

##### Gene expression analysis via qPCR

To examine gene expression, qPCR was performed on the Thermal Cycler CFX384 Touch Real-Time PCR Detection System (C1000 Touch, BIO-RAD). RNA was isolated with the ReliaPrep TM RNA Tissue Miniprep System (Promega) following the manufacturer's instructions. 200 – 300 ng RNA were transcribed into cDNA using the iScript cDNA Kit (BioRad) and subsequently diluted to a concentration of 5 ng/µL. Each cDNA sample was analyzed in duplicates. 11 µL master mix and 1 µL cDNA sample were used per well (Suppl.Tab. 2) with the following settings: Initial denaturation at 95 °C for 3 min, followed by 40 cycles of denaturation at 95 °C for 10 sec, annealing at 60 °C for 20 sec, elongation at 72 °C for 30 sec and final denaturation 60 °C for 5 sec followed applying a melt curve from 60-95 °C (5 sec). Suppl. Tab. 3 summarizes the primers used.

***Suppl.Tab. 2: Master Mix components for qPCR.***

| Master Mix component | Volume |
|---|---|
| IQ SYBR Green Supermix (BioRad) | 6.0 µL |
| Primer (stock of 10 mM containing both forward and reverse) | 0.6 µL |
| Nuclease-free water | 4.4 µL |
| cDNA (5 ng/µL) | 1 µL |

*Suppl.Tab. 3: Primers used in qPCR. Annealing temperature is 60 °C for all primer pairs.*

| Gene | Forward 5´- 3´ | Reverse 5´- 3´ |
|---|---|---|
| CPT1B | TGTATCGCCGTAAACTGGACCG | TGTCTGAGAGGTGCTGTAGCAC |
| GAPDH | GTC TCC TCT GAC TTC AAC AGC G | ACC ACC CTG TTG CTG TAG CCA A |
| HPRT | CATTATGCTGAGGATTTGGAAAGG | CTTGAGCACACAGAGGGCTACA |
| KCNJ2 | AACAGTGCAGGAGCCGCTTTGT | AGGACGAAAGCCAGGCAGAAGA |
| MYH7 | GGAGTTCACACGCCTCAAAGAG | TCCTCAGCATCTGCCAGGTTGT |
| MYL2 | CGGAGAGGTTTTCCAAGGAGGA | CTCTTCTCCGTGGGTGATGATG |
| NR2F2 | CCGACCGGGTGGTCGCCTTTATGGA | CGGCTGGTTGGGGTACTGGCTCCTA |
| PDK4 | AGGTGGAGCATTTCTCGCGCTA | GAATGTTGGCGAGTCTCACAGG |
| SCN5a | TACACCTTTGAGTCTCTGGTCAAG | AATCACACTAAAGTCCAGCCAGTT |
| TTN_N2B | CCA ATG AGT ATG GCA GTG TCA | TAC GTT CCG GAA GTA ATT TGC |

**Immunofluorescence stainings**

For immunofluorescence, the cells were fixed in 4% Histofix (Roth). For 2D cultured cells and dissociated spheroids cells were fixed 20 min in 4% Histofix at room temperature, subsequently washed in 1xPBS (#14190-144, Gibco) and stored in Blocking buffer (1% BSA/PBS V/V). The cover slips were washed thrice in 1xPBS at room temperature and subsequently incubated in Staining Buffer 1 (Blocking solution supplemented with 0.1% Triton-X) with the primary antibody overnight at 4 °C. The next day, the cover slip was washed thrice with 1xPBS followed by incubation with secondary antibody in Staining Buffer 1 for 1,5 h at room temperature in the dark. Afterwards the coverslip was washed again in 1xPBS and mounted with Fluoromount-G (Southern Biotech). In a subset of samples, 250 nM Mitospy-orange (Biolegend) was added for 30 min before fixation. Sarcomere and Mitospy imaging were performed using a widefield microscope (Leica THUNDER) with a 63x oil immersion objective. Analysis of circularity and sarcomeric length was analyzed in Image J using the circularity index and sarcOptim, respectively. The mitochondrial network was quantified using the Image J plugin MiNa.

For whole spheroid stainings (not mounted), fixed spheroids were permeabilized overnight at 4 °C on a shaker using a permeabilization solution (PBS supplemented with 2.5% Normal Donkey Serum [NDS], 5% Bovine Serum Albumin [BSA] and 0.25% Triton X-100, V/V). After permeabilization, the spheroids were incubated with primary antibodies for 72 h at 4 °C: anti-sarcomeric alpha-actinin (rabbit, 1:200) and anti-CD31 (mouse, 1:200). These antibodies were diluted in an antibody diluent solution (PBS supplemented with 2.5% NDS, 1% BSA and 0.01% Triton X-100, V/V). The samples were gently washed three times for 30 min each with PBST (PBS with 0.05% Tween 20). Following the washing steps, the spheroids were incubated with secondary antibodies for 72 h at 4 °C: donkey anti-rabbit-Alexa Fluor 647 (1:400) and donkey anti-mouse-Alexa Fluor 546 (1:400), diluted in antibody diluent solution. After incubation with secondary antibodies, the organoids were washed three times for 30 min with PBST. They were then incubated with the Hoechst 33342 nucleic acid stain (1:400) in PBS for 2 h at 4°C. Finally, the samples were washed three times for 5 min, and then stored at 4°C. Whole spheroid imaging was performed in glass-bottom microscopy dishes, using an inverted confocal microscope (Leica TCS SP8 X, excitation light provided by a 405-nm diode laser and selected lines of a white light laser, fluorescence recorded with a photon multiplier [Hoechst] or sensitive hybrid detectors operated in photon-counting mode, using a 40x water immersion objective or a 63x glycerol immersion objective).

***Suppl.Tab. 4: Antibodies and antibody dilutions used in this study.***

| Antibody | Company | Dilution |
|---|---|---|
| ***Primary antibodies*** | | |
| MLC2v (2D) | Proteintech #10906-1-AP | 1:200 |
| MLC2a (2D) | Synaptic Systems #311-011 | 1:200 |
| Titin-M (2D) | Myomedix-order form | 1:500 |
| CD31 (2D) | Thermo fisher #MA1-26196 | 1:200 |
| CD31 (spheroid) | Thermo Fisher Scientific #MA5-13188 | 1:200 |
| α-actinin (spheroid) | Abcam, ab137346 | 1:200 |
| ***Secondary antibodies*** | | |
| Alexa488-anti mouse | Life technologies #A10680 | 1:500 |
| Alexa555-anti rabbit | Life technologies #A31572 | 1:500 |
| Alexa647-anti rabbit | Life technologies #A31573 | 1:500 |
| Alexa647-anti rabbit (spheroid) | Thermo Fisher Scientific #A21123 | 1:400 |
| Alexa546-anti mouse (spheroid) | Thermo Fisher Scientific #A21123 | 1:400 |
| ***Mitochondrial dye*** | | |
| Mitospy-orange | BioLegend, 250 nM for 30 min at RT in culture medium before fixation | |
| ***Nuclear stain*** | | |
| Hoechst 33342 (spheroids) | Abcam #ab228551 | 1:400 |

#### Supplemental Figures:

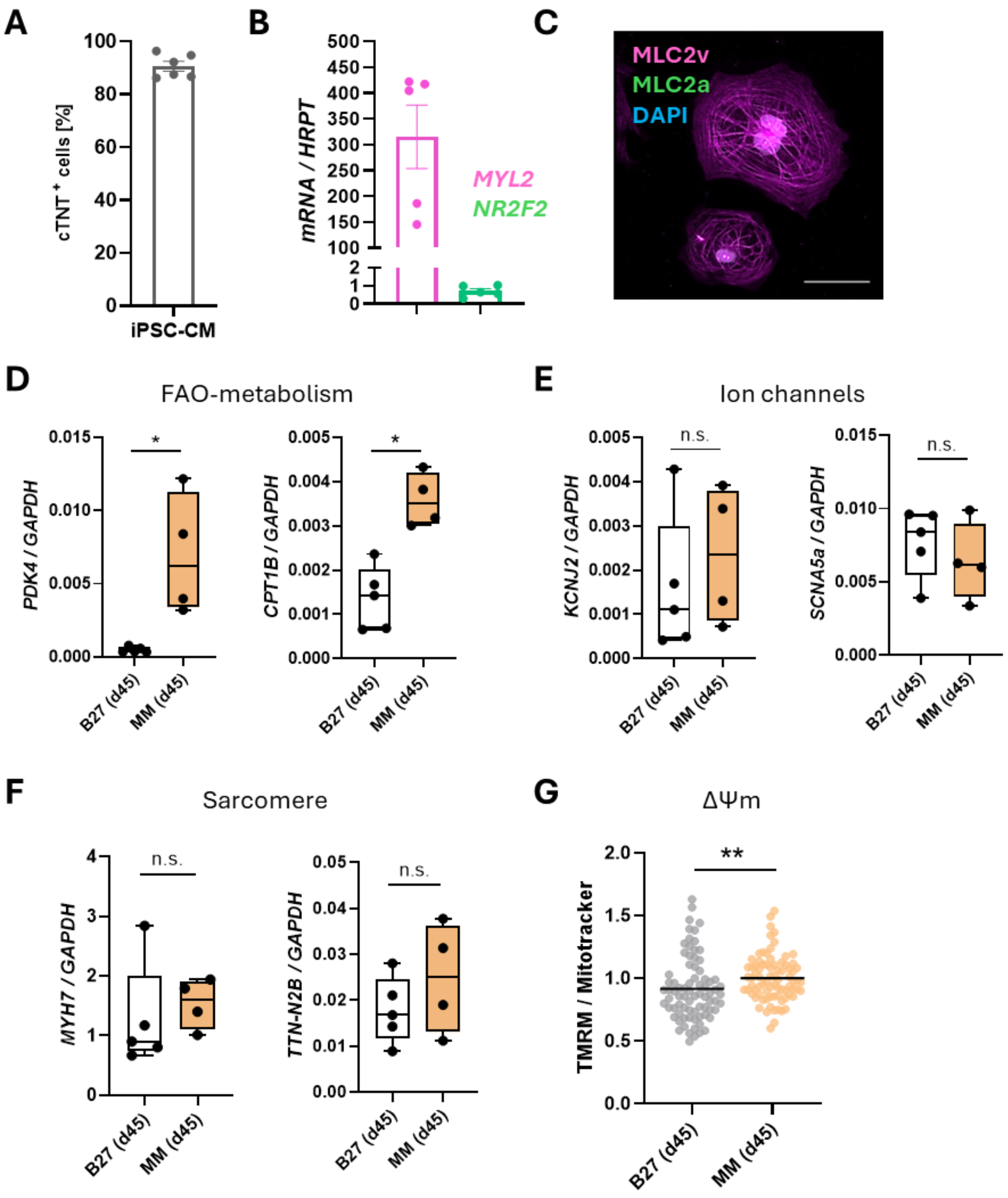


**Suppl. Fig. 1: Quality assessment and effects of the novel-composite maturation medium (MM).**

**A:** Cells were stained with the cardiac marker cTNT and analyzed using Flow cytometry to determine the % of cTNT-positive cells (2D culture). Each dot represents one cardiac differentiation.

**B:** QPCR analysis of the ventricular marker MYL2 and the atrial marker NR2F2. HPRT serves for normalization (2D culture). Each dot represents one cardiac differentiation.

**C:** Immunofluorescence staining of iPSC-CM (2D culture); antibody against MLC2v (pink) as (ventricular marker) and MLC2a (green) as atrial marker, and DAPI (cyan) to stain the nucleus. Scale bar: 50 µm.

**D-E:** QPCR analysis of different genes involved in fatty-acid oxidation (FAO) (D), expressing ion channels (E) or sarcomere proteins (F). HPRT serves for normalization. Each dot represents one cardiac differentiation. Statistical testing by Mann-Whitney test.
**G:** Quantification of mitochondrial membrane potential ($\Delta\Psi m$). [Number of differentiations/analyzed cells] for B27(d45) [4/78] and MM(d45) [4/80]. Statistical testing by Mann-Whitney test.

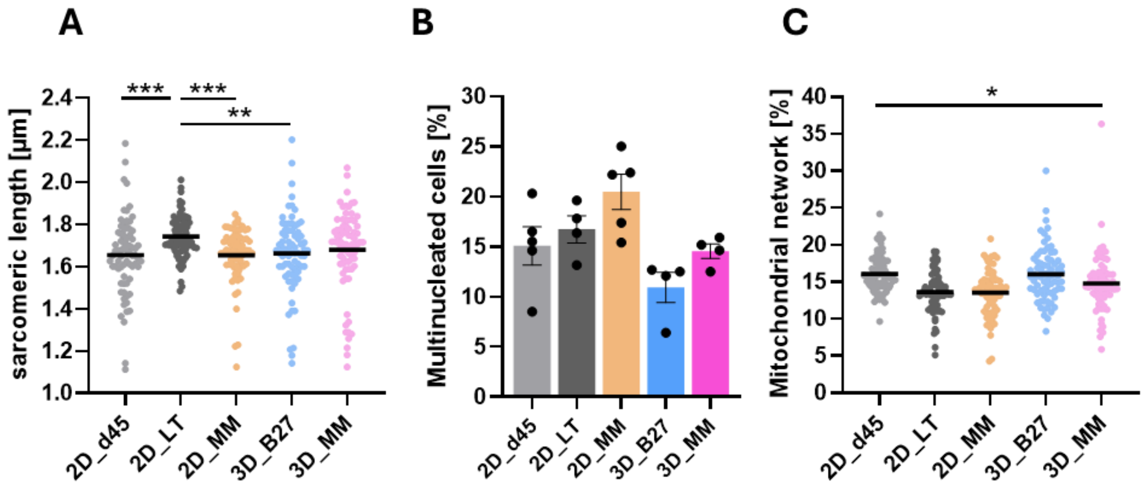


**Suppl. Fig. 2: Analysis of sarcomeric length, nucleus numbers and the mitochondrial architecture.**

**A:** Quantification of sarcomere length. The titin staining was used to determine the length using the SarcOptim plugin in Image J. Data is represented as beeswarm blot with mean. [No of differentiations / analyzed pictures] for 2D_d45 [4/73], 2D_LT [4/85], 2D_MM [4/71], 3D_B27 [4/74] and 3D_MM [4/78]. P-values were calculated by the Kruskal-Wallis test with Dunn´s multiple comparisons test. Significant values are marked * p<0.05 , ** p<0.01, ***p<0.001.

**B:** Percentage of multinucleated cells: Total number of CM divided by number of CM with more than one nucleus. Images with DAPI for nucleus and titin-M for cardiomyocyte marker were used for manual counting. Data is represented as mean with SEM. Each dot represents one cardiac differentiation. Statistical analysis with Kruskal-Wallis test against 3D_MM with Dunn´s correction. No significances were detected.

**C:** Analysis of mitochondrial network based on Mitospy stainings and image analysis with the MiNa plugin in ImageJ. Data is represented as beeswarm blot with mean. [No of differentiations / analyzed pictures] for 2D_d45 [4/73], 2D_LT [4/65], 2D_MM [4/71], 3D_B27 [4/74] and 3D_MM [4/72]. Statistical analysis with Kruskal-Wallis test against 3D_MM with Dunn´s correction. Significant values are marked * p<0.05 , ** p<0.01, ***p<0.001.

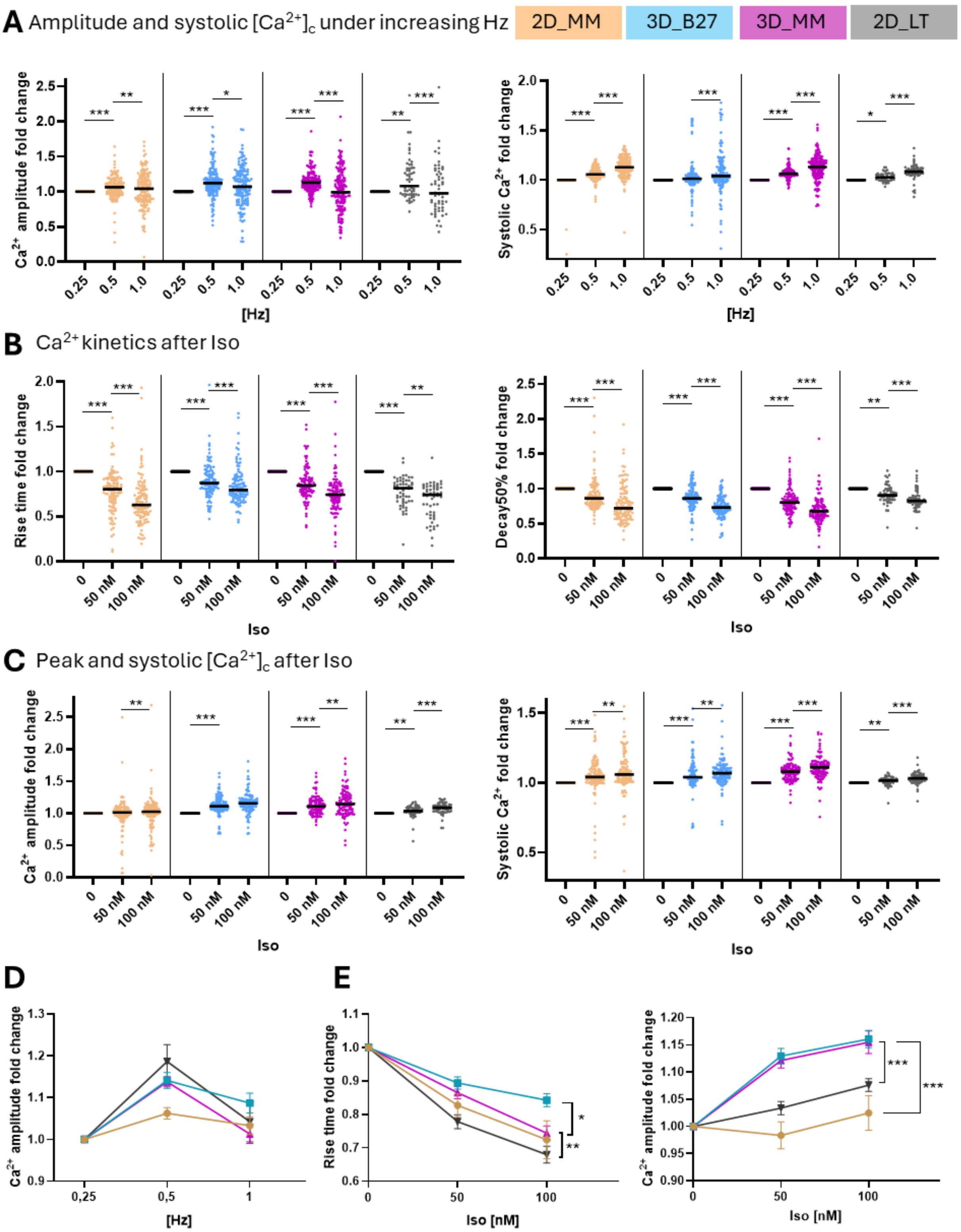


**Suppl. Fig. 3: $Ca^{2+}$ handling parameters after electrical or Iso-treated stimulation.**
Measurement of $Ca^{2+}$ cycling with ratiometric Indo-1 dye (data as ratio 405/495).
**A:** The iPSC-CM from different maturation conditions were subjected to increasing pacing (0.25 Hz to 0.5 Hz to 1Hz). Since matching cells were measured the basal condition is set to 1 and data is shown as fold change. Statistics was performed by Friedmans test´with Dunn´s multiple comparisons slected for 0.25 Hz vs 0.5 Hz and 0.5 Hz vs 1.0 Hz within each maturation cohort.

**B-C:** The iPSC-CM from different maturation conditions were subjected to increasing Iso concentrations (0 nM to 50 nM to 100 nM) at 0.5 Hz pacing. Analysis of **B:** $Ca^{2+}$ kinetics parameter rise and dacay50 time and **C:** the peak height and systolic $Ca^{2+}$. Since matching cells were measured the basal condition is set to 1 and data is shown as fold change of the respective $Ca^{2+}$ parameter. Statistics was performed by Friedmans test´with Dunn´s multiple comparisons slected for 0 nM vs 50 nM and 50 nM vs 100 nM within each maturation cohort.
**D+E:** The iPSC-CM from different maturation conditions were subjected to increasing pacing **(D)** or Iso concentrations **(E)**. Since matching cells were measured the basal condition is set to 1 and data is shown as fold change of the respective $Ca^{2+}$ parameter. Statistics was performed by Two-way ANOVA with Geisser-Greenhouse´s correction. P-value (column factor) for 3D_MM vs 2D_MM; 3D_MM vs 3D_B27 and 3D_MM vs 2D_LT.
**A-E:** [No. of differentiations / analyzed cells] for 2D_MM [5/145], 3D_B27 [5/137], 3D_MM [5/137] and 2D_LT [2/57]. Significant values are marked * $p<0.05$ , ** $p<0.01$, ***$p<0.001$. Iso = Isoprenaline
